# Model Agnostic Semi-Supervised Meta-Learning Elucidates Understudied Out-of-distribution Molecular Interactions

**DOI:** 10.1101/2023.05.17.541172

**Authors:** You Wu, Li Xie, Yang Liu, Lei Xie

## Abstract

Many biological problems are understudied due to experimental limitations and human biases. Although deep learning is promising in accelerating scientific discovery, its power compromises when applied to problems with scarcely labeled data and data distribution shifts. We developed a semi-supervised meta learning framework Meta Model Agnostic Pseudo Label Learning (MMAPLE) to address these challenges by effectively exploring out-of-distribution (OOD) unlabeled data when transfer learning fails. The power of MMAPLE is demonstrated in multiple applications: predicting OOD drug-target interactions, hidden human metabolite-enzyme interactions, and understudied interspecies microbiome metabolite-human receptor interactions, where chemicals or proteins in unseen data are dramatically different from those in training data. MMAPLE achieves 11% to 242% improvement in the prediction-recall on multiple OOD benchmarks over baseline models. Using MMAPLE, we reveal novel interspecies metaboliteprotein interactions that are validated by bioactivity assays and fill in missing links in microbiome-human interactions. MMAPLE is a general framework to explore previously unrecognized biological domains beyond the reach of present experimental and computational techniques.

## 1 Introduction

Small molecule-protein interactions within an organism or across species influence diverse biological processes. However, a majority of molecular interactions are understudied due to experimental limitations and human biases. Unraveling these unrecognized molecular interactions is essential for gaining critical insights into the secrets of life and is instrumental in the field of drug discovery.

Metabolite-protein interactions (MPIs) play a crucial role in regulating metabolic pathways, triggering signaling pathways, and maintaining cellular homeostasis.

However, MPIs are frequently low-affinity and are difficult to be detected by experiments. A recent study discovered that many overlooked MPIs contribute to the survival and growth of organisms in response to a changing environment[1]. Additionally, proteome-wide characterization of MPIs provides strong evidence that metabolites serve as not only intermediates in metabolic reactions but also signaling molecules via interactions with proteins in which the metabolite is not a substrate for metabolic reactions[2][3]. Furthermore, the microbiome coevolves with the human and plays a part in shaping human phenotypes[4]. In the human body, microbiota produces an extremely diverse metabolite repertoire that can gain access to and interact with host cells, thus influencing the phenotype of the human host[5]. For example, butyrate produced by the microbiome binds to human non-enzyme receptors such as GPCRs[6]. The human microbiome is not only associated with a large number of human diseases but also responsible for the efficacy and toxicity of therapeutics (e.g. cancer immunotherapy) that target human host[7][8]. Thus, elucidating previously unrecognized intraspecies and interspecies MPIs will uncover the fundamental rule of life and revolutionize biomedicine[9].

Uncovering novel drug-target interactions (DTI) will facilitate identifying novel therapeutic targets, understanding polypharmacology, and advancing drug repurposing, thereby accelerating drug discovery and development. Unfortunately, small molecule ligands of more than 90% protein families remain unknown[10].

Many of these understudied proteins could be potential drug targets[11]. The lack of knowledge of endogenous and extraneous ligands of understudied proteins hinders the drug development of presently incurable diseases[12]. On the one hand, many disease-causing genes are functionally and pharmaceutically uncharacterized[13]. It is a time-consuming and high-risk endeavor to develop assays for compound screening of understudied proteins. On the other hand, a drug acts on a biological system, and often interacts with not only its intended target but also unknown off-target(s) that may lead to unexpected side effects[14]. Thus, identifying drug-target interactions including chemicals with novel scaffolds will improve the successful rate of drug discovery for unmet medical needs[15].

Given the experimental challenges in elucidating understudied molecular interactions, deep learning offers a promising alternative approach owning to its recent phenomenal successes in Natural Language Processing (NLP) and image processing. A number of deep learning models have been developed to predict drug-target interactions[16, 17, 18]. Nevertheless, few methods can accurately and reliably predict understudied molecular interactions due to the dearth of labeled data and out-of-distribution (OOD) problems in which small molecules or proteins involved in the interaction are significantly different from those in the annotated databases used as training data. A plethora of transfer learning techniques have been developed to address the molecular OOD problem[19]. However, they offer little help in bridging remote chemical spaces[19]. Recently, semi-supervised learning and meta-learning alone have shown promise in addressing OOD challenges in protein-ligand interaction predictions[20, 10]. Semi-supervising learning uses the labeled data to learn and then generalizes that knowledge to the unlabeled data. It is a potentially powerful technique to explore uncharted protein and chemical spaces. Meta-learning is an approach to learning to learn. It has demonstrated superior generalization power in many applications[21, 10, 22]. However, to our knowledge, few works have combined these two techniques to address the data scarcity and OOD challenges in molecular interaction predictions.

In this paper, we have developed MMAPLE Meta Model Agnostic Pseudo Label Learning to address the challenges aforementioned. MMAPLE is effective in exploring unlabeled data and addressing the OOD problem. We have demonstrated that MMAPLE significantly improves the accuracy of OOD DTI, hidden human metabolite-enzyme interactions, and understudied microbiome-human MPI predictions on multiple baseline models. Using MMAPLE, we have identified and experimentally validated novel microbiome-human MPIs and proposed their associations with human biology. Our findings suggest that MMAPLE can be a general framework for investigating understudied biological problems.

## 2 Overview of MMAPLE

We evaluate the proposed MMAPLE method on three diverse cases: OOD DTIs, hidden human MPIs, and understudied microbiome-human MPIs. The statistics of training/validation and testing data are shown in Figure1(A). The distribution of chemical similarities between training/validation and testing data is shown in Supplemental Figure S1. In brief, for OOD DTIs and hidden human MPIs, no chemicals in the testing data have a Tanimoto coefficient larger than 0.5 compared with those in the training/validation set. Although 1.7% chemicals in the testing data are similar to those in the training/validation data of microbiome-human MPIs, the proteins are significantly different based on Evalue as shown in Supplemental Figure S1(D). Furthermore, there is no labeled microbiome-human MPIs in the training data. Thus, all benchmarks are in a challenging OOD or understudied label scarcity scenario.

The rationale MMAPLE for predicting understudied or OOD molecular interactions is to iteratively transfer the knowledge of observed molecular interactions to the unexplored chemical genomics space of interest (Figure1(B)). Specifically, we train base models that used labeled molecular interactions from ChEMBL[23]. The base models used in this study included four state-of-theart models for chemical-protein predictions: pre-trained protein language model DISAE[24], TransformerCPI[16], DeepPurpose[17], and BACPI[18]. Then MMAPLE was applied to these models for exploring unlabeled molecular interaction space, as illustrated in Figure 1(B). MMAPLE first initializes a teacher model using the labeled data. Then a sampling strategy is applied to select a set of unlabeled data from the large space of understudied OOD DTIs or MPIs. The pre-trained teacher model makes predictions about the selected unlabeled data and assigns labels to them (pseudo labels). Next, a student model is trained using pseudolabeled data. Different from conventional teacher-student model, the student model is evaluated by labeled data and provides feedback (metadata) to the teacher model during training the student model. Finally, the teacher model is updated based on the performance of the student and generates new pseudo labels. This process repeats multiple times. The details of MMAPLE are in the Method section.

**Figure 1:**
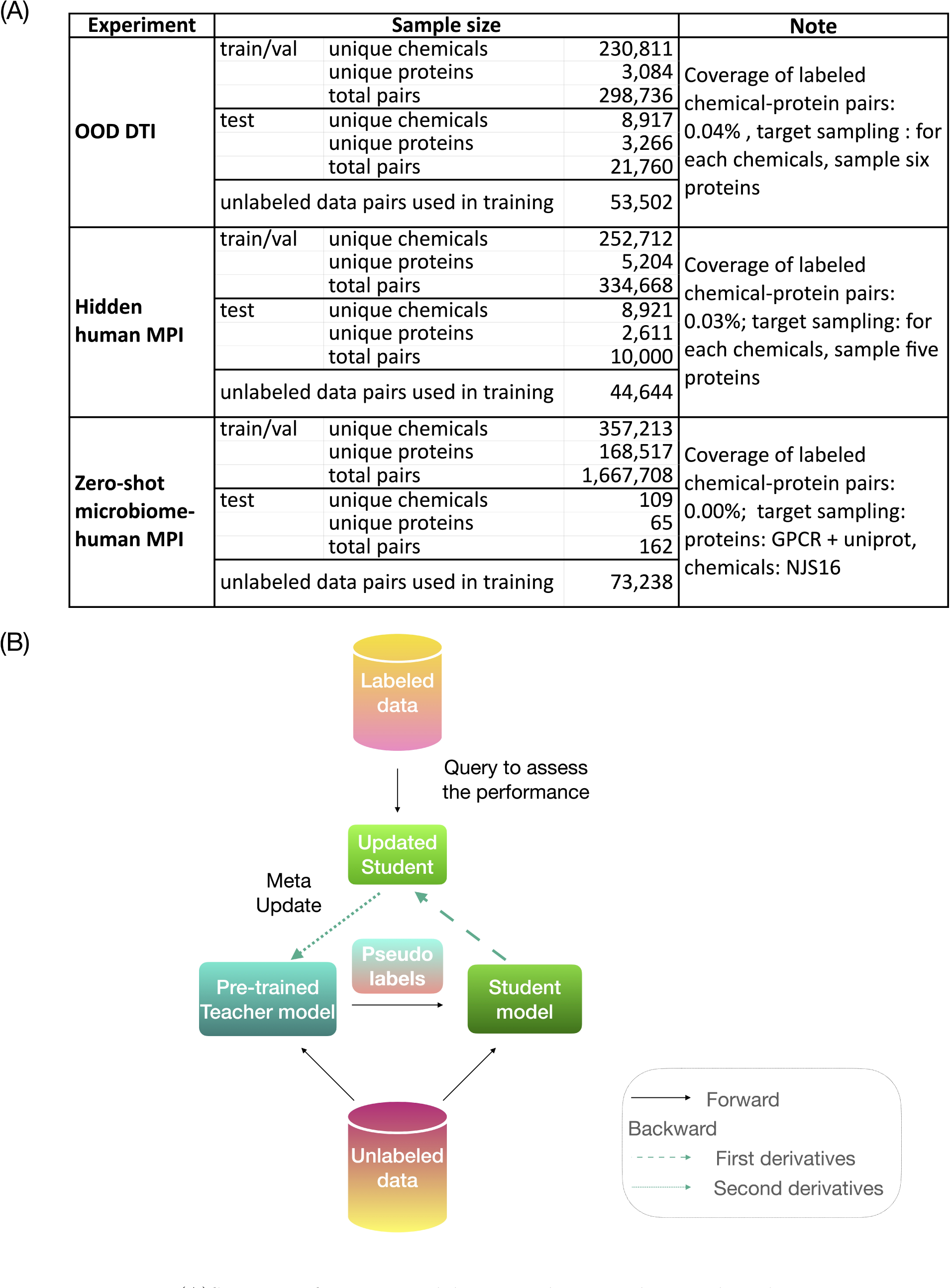
(A)Statistics of training, validation, and testing data used in this study. (B)Illustration of MMAPLE framework. A deep learning model is trained using both labeled and unlabeled data_2_a_2_nd iteratively updated using gradients from the trained model as metadata.

## 3 MMAPLE significantly improves the performance of OOD DTI predictions

We first evaluated the performance of MMAPLE for OOD DTI predictions. We used molecular interactions in ChEMBL[23] and HMDB[25] as training data, and annotated DTIs from DrugBank[26] as testing data. To simulate an OOD scenario, we removed all chemicals that are structurally similar to drugs in the testing data (Tanimoto coefficient *>* 0.5). As shown in Figure 2, both PRAUCs and ROC-AUCs of MMAPLE are significantly improved over all baseline models with p-values less than 0.05 (Supplemental Table S1,S2). The percentage of improvement on PR-AUC ranges from 13% to 26%. Furthermore, the trained models are less over-fitted than the baseline models, as supported by the narrow gaps between the training curve of validation data and that of testing data, shown in Supplemental Figure S2.

**Figure 2:**
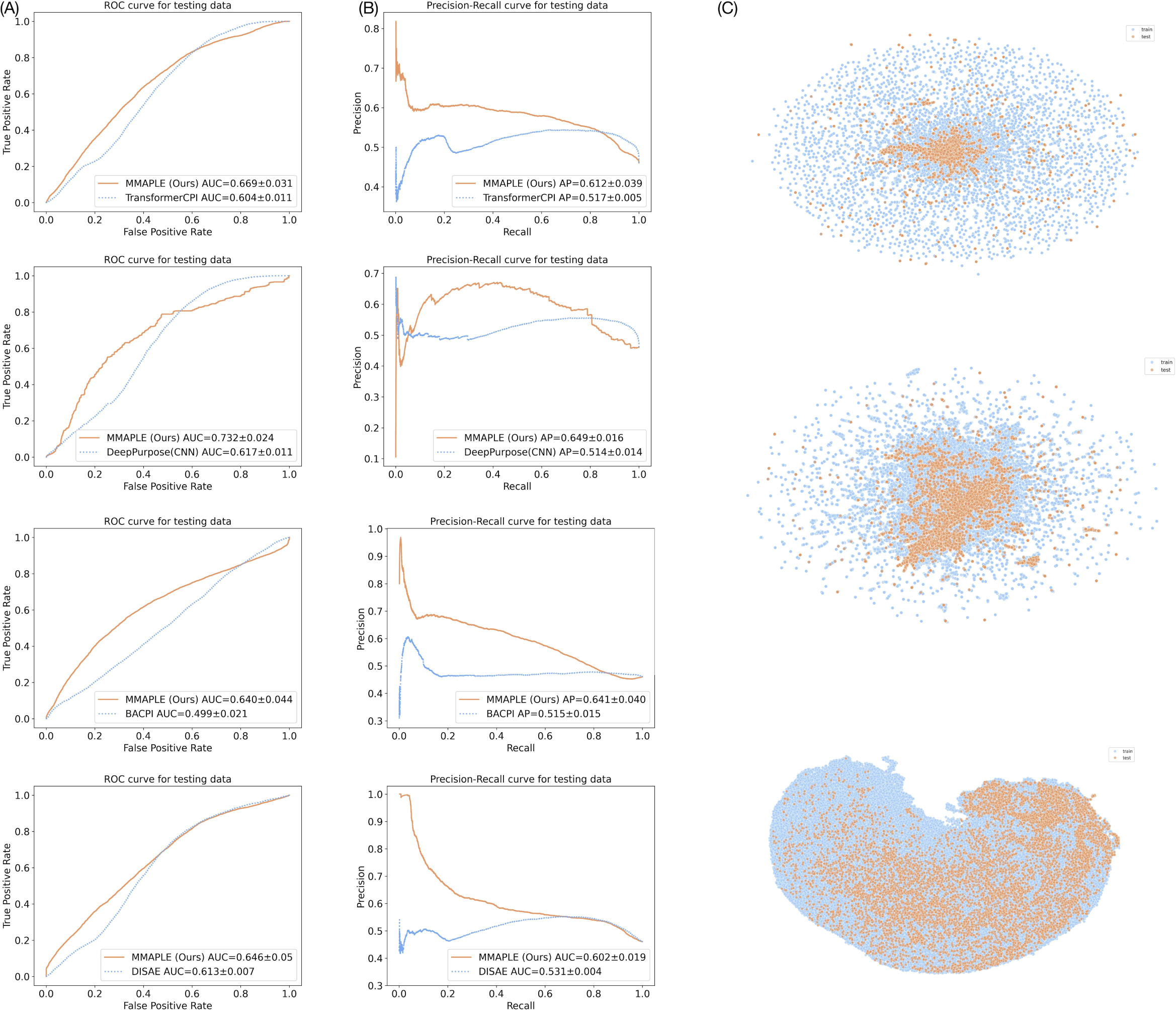
OOD DTI prediction outcomes when applying MMAPLE to baseline models. (A) ROC curves. (B) PR curves. (C) UMAP visualization of chemical space. top to bottom original fingerprints, baseline embeddings from DISAE, and MMAPLE embeddings. Orange and blue dots for MMAPLE and baseline, respectively.

**Figure 3:**
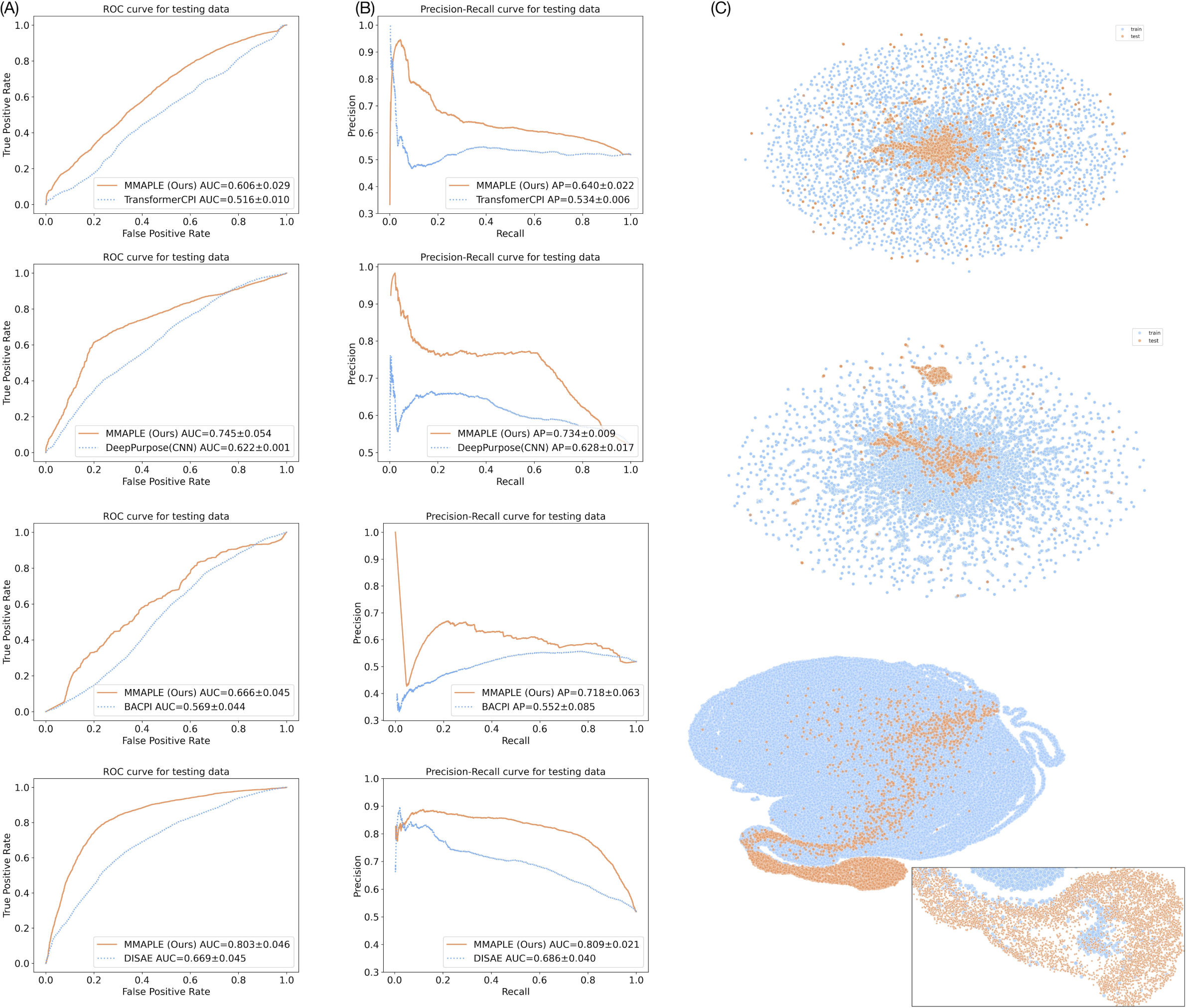
Hidden Human MPI prediction outcomes when applying MMAPLE to baseline models. (A) ROC curves; (B) PR curves; (C) UMAP visualization of chemical space. top to bottom original fingerprints, baseline embeddings from DISAE, and MMAPLE embeddings. Orange and blue dots for MMAPLE and baseline, respectively.

The superior performance of MMAPLE may be because it can better align the embedding space of OOD samples to that of training data. To test this hypothesis, we investigated if MMAPLE could alleviate the distribution shift between training and testing data. We extracted the embeddings of the training and testing examples before training and acquired by DISAE and MMAPLE, then utilized the Uniform Manifold Approximation and Projection (UMAP) for visual analysis. Figure2(C) supports this hypothesis. Before the training, the embeddings of training chemicals are scattered around those of testing chemicals. While DISAE the best-performed baseline model narrows the dispersion, our model achieves tighter overlap between two distributions. Importantly, our model not only draws them closer but also ensures a more uniform distribution within each group, reducing inter-distribution gaps.

Transfer learning in the protein space via protein language modeling can improve the performance of DTI prediction[24]. As shown in Figure2(A, B) and Figure3(A, B) in the next section, DISAE that is based on a pre-trained protein language model outperforms other baselines that do not utilize the language model. However, the improvement from DISAE is not as significant as that by MMAPLE. Additionally, We study if transfer learning in the chemical space could boost the performance of OOD DTI predictions. We apply a chemical pretraining-fine-tuning based on self-supervised Motif Learning Graph Neural Network (MoLGNN)[27]. In consistent with recent findings[19], no improvement was detected as shown in Supplemental Figure S3.

## 4 MMAPLE significantly improves the performance of hidden human MPI predictions

We next evaluated the performance of the MMAPLE model in predicting hidden human MPIs. We first trained the model using ChEMBL which primarily includes exogenous small molecule ligands and druggable protein targets. We evaluated the performance of the trained model using the Human metabolite database HMDB[25] on human MPIs. The test cases were in the OOD setting, as supported by the chemical similarity distributions (Supplemental Figure S1). Figure3(A)indicates that MMAPLE significantly outperforms all of state-ofthe-art baseline models on both ROC and PR. The ROC-AUC and PR-AUC increase by 17% to 20% and 17% to 30%, respectively, suggesting that MMAPLE is able to accurately predict hidden human MPIs in an OOD setting.

Again, MMAPLE training brings the embeddings of testing samples closer to those of training data than the baseline, as shown in Figure3(C). Overall, our results suggest that MMAPLE significantly outperforms the state-of-theart methods for OOD DTI and hidden MPI predictions.

## 5 MMAPLE significantly improves the performance of understudied interspecies MPI pre-dictions and reveals the molecular basis of microbiomehuman interactions

Known interspecies microbiome-human MPIs are extremely scarce, only including 17 observed active interactions (See Methods for details). To investigate interspecies interaction, MMAPLE was trained on a combination of three datasets: HMDB, ChEMBL, and NJS16[28], while the test set consisted of 17 annotated along with 145 negative microbiome-human MPIs from the literature[29][30]. As shown in Figure 1(A) and Supplemental Figure S1, No metabolite-protein pairs in the testing set have similar chemicals or proteins to those in the training/validation set. Because no interspecies MPIs exist in the training and validation set, the problem is a zero-shot learning scenario. The previously best performed model DISAE has a poor performance, with a PR-AUC of 0.193. It indicates that transfer learning alone is not sufficient to address the OOD challenge of interspecies MPI predictions. Our results, presented in Figure 4(A), demonstrate that MMAPLE significantly outperforms DISAE in terms of ROC and PR. It achieves a three-fold increase in the PR-AUC for interspecies MPI predictions. These findings indicate that MMAPLE holds promise in deepening our comprehension of interspecies interactions, thus serving as a valuable tool for investigating the impact of the microbiome on human health and disease.

**Figure 4:**
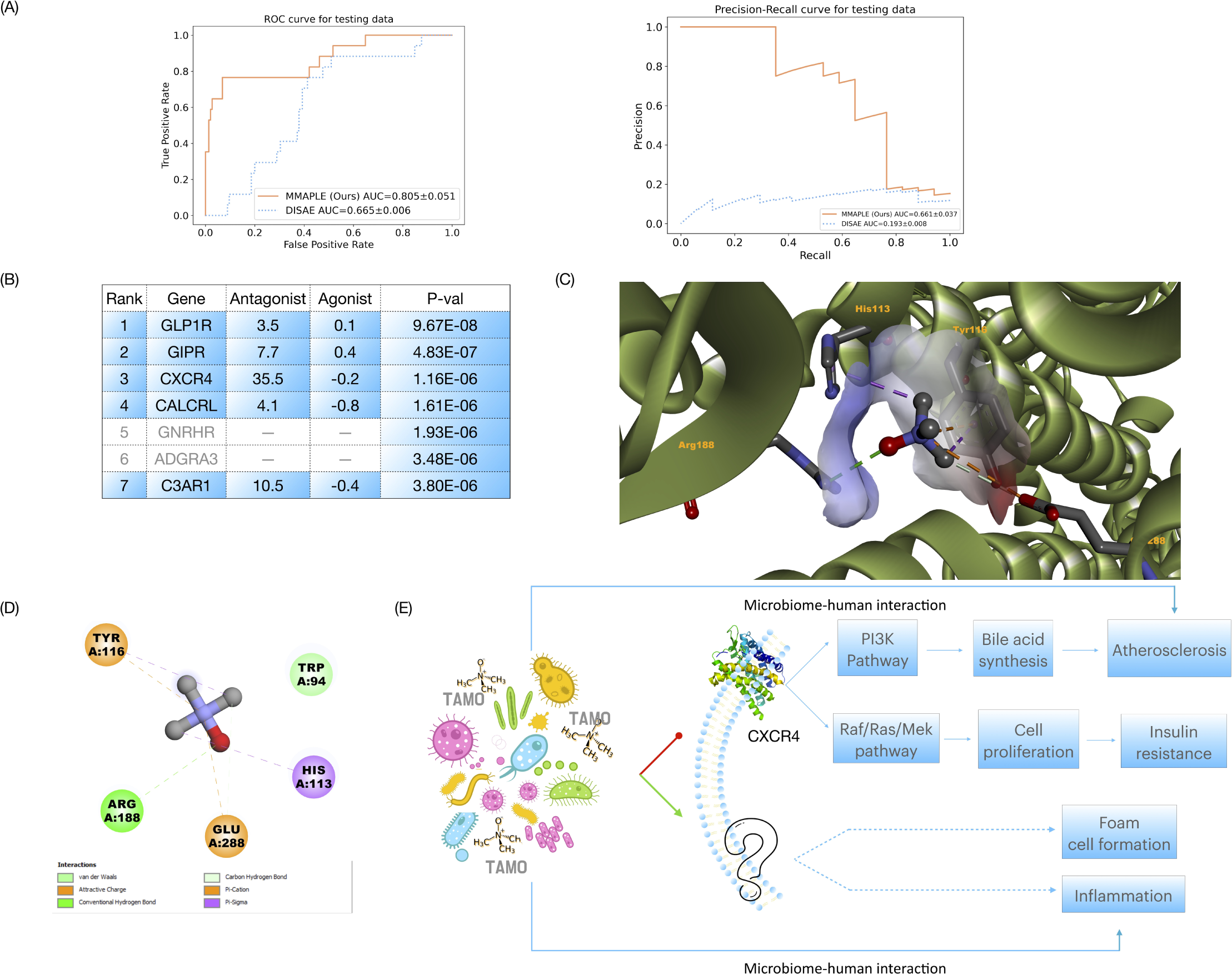
Results of TAMO-GPCR interactions prediction when applying MMAPLE to DISAE. (A) ROC and PR curves; (B) Top 7 predicted G-protein coupled receptor (GPCR) genes that interact with TAMO and GPCR functional assay results, with p-value indicating the tail probability from Kernel density estimation; (C) Predicted 3D binding pose of TAMO on the CXCR4 antagonist conformation; (D) Interaction patterns between TAMO and CXCR4; (E) Proposed molecular mechanism of TAMO-human interactions. No assay is available for GNRHR and ADGRA3.

To further validate the performance of MMAPLE, we predicted and experimentally validated the interactions between trimethylamine N-oxide (TAMO) and human G-protein coupled receptors (GPCRs). TAMO is a small molecule generated by gut microbial metabolism. It has been observed that elevated plasma levels of TMAO increase the risk for major adverse cardiovascular events[31], activate inflammatory pathways[32], and promote foam cell formation[33]. Additionally, TMAO inhibits insulin signaling[34]. However, it remains elusive how TAMO modulates these pathological processes at a molecular level. Besides its biological interest, TAMO is one of the most challenging molecules for MMAPLE. Firstly, the current study of microbiome-human interactions mainly focuses on short-chain fatty acids, there are few data for TAMO. Secondly, TAMO is a molecule with different structural characteristics from other chemicals in the training data. Thus, we choose TAMO to rigorously evaluate MMAPLE in an OOD scenario.

Figure 4(B) lists the top 7 predicted GPCR genes that interact with TAMO with a p-value less than 5.0e-5. We performed GPCR functional assays to experimentally test the binding activities of five of them under the concentration of 30 *µM* of TAMO, which is the same concentration used in the previous study and is based on the physiological concentration of TAMO in the human (1-45 *µM*)[35]. The assay for two top-ranked GPCRs CNRHR and ADGRA3 is not available. As shown in Figure 4(B), all five tested GPCRs show antagonist activities, and CXCR4 demonstrates a strong activity (activity score*>*30). The full predictive results can be found in Supplemental Table S3.

Protein-ligand docking suggested that TAMO can fit into the antagonist conformation of the CXCR4 structure, as shown in Figure 4(C) and (D). AutoDock Vina [36] was applied on TAMO to find the best conformation in the CXCR4 chemokine receptor (PDB ID: 3ODU). The docking conformations with the best (lowest) predicted binding energies were selected and the interactions between TAMO and 3ODU were shown in Figure 4(C) and (D). Among these interacting residues, TRP94, TYR116, and Glu288 also interact with the co-crystallized ligand of 3ODU. TYR116 and GLU288 provide attractive charges to the nitrogen atom on TAMO. ARG188 forms a conventional hydrogen bond with an oxygen atom on TAMO. These strong interactions could keep TAMO in the binding pocket.

The CXCR4 antagonism by TAMO establishes a causal linkage for observed microbiome TAMO-human interactions, as illustrated in Figure 4(E). It is known that CXCR4 regulates PI3K and RAF/RAS/MEK pathways [37] (KEGG Pathway: https://www.genome.jp/pathway/hsa04062). PI3K pathway regulates bile acid synthesis[37]. TAMO’s inhibition on bile acid synthesis may be responsible for its promotion effect on atherosclerosis[31]. The physiological effect of TAMO on obesity and insulin resistance may be via CXCR4-RAF/RAS/MEK axis. It has been observed that the deficiency of CXCR4 and impaired RAF/RAS/MEK signaling results in obesity and insulin resistance[38][39][40][41]. Additionally, CXCR4 is important for cell formation via the RAF/RAS/MEK pathway[42]. Thus, microbiome TAMO-human CXCR4 interaction is responsible for the several observed pathological effects of TAMO. However, several other TAMO effects such as foam cell formation and inflammation cannot be directly explained by the TAMO-CXCR4 interaction. It is possible that other human proteins can interact with TAMO.

## 6 Semi-supervised learning and meta-learning synergistically contribute to the performance of MMAPLE

In our comprehensive ablation study, we rigorously examined the influence of several key components on our model’s performance: the introduction of target sampling and semi-supervised learning with pseudo labels, the choice between utilizing soft pseudo labels or hard labels, and the application of meta-learning. When excluding meta-learning from our training process, we kept the teacher model static, therefore restricting it to generate constant pseudo labels for the student model to learn from. This leads to performance decline when compared to our full MMAPLE model, as shown in Table1. The absence of iterative feedback learning addresses the critical role of meta-learning.

To investigate the effect of pseudo labels, we trained the model sorely with meta-learning by leveraging the Model-Agnostic-Meta-Learning (MAML) framework[43]. This approach, while constantly outperforming the baseline, resulted in 124 % fall on PR-AUC compared to MMAPLE. This experiment not only demonstrated the intricate dependencies between meta-learning and semi-supervised learning but also underscored the necessity of synergy of these techniques to achieve superior model performance.

Models trained on one-hot (hard) labels are subject to over-fitting since they do not represent soft decision boundaries across concepts. Soft labels, which are probability distributions over the possible classes as opposed to hard labels, are often demonstrated to be more effective due to the ability to provide the model with more information about the uncertainty in the data, as well as the ability against label noise, resulting in more robust predictions[44][45]. As shown in the table 1, when soft labels were used, ROC-AUC improved by 25%, and PR-AUC increased by twofold.

**Table 1:** Ablation study.

| Methods | ROC-AUC | PR-AUC |
| --- | --- | --- |
| w/o MMAPLE (baseline) | $0.665 \pm 0.006$ | $0.193 \pm 0.008$ |
| w/o target sampling | $0.727 \pm 0.033$ | $0.308 \pm 0.020$ |
| w/o soft labels | $0.708 \pm 0.013$ | $0.247 \pm 0.024$ |
| w/o pseudo labels | $0.748 \pm 0.009$ | $0.295 \pm 0.034$ |
| w/o meta-learning | $0.791 \pm 0.061$ | $0.632 \pm 0.108$ |
| MMAPLE | $0.805 \pm 0.051$ | $0.661 \pm 0.037$ |

Target sampling is a widely used technique in semi-supervised learning to increase the performance of a model by generating pseudo labels for unlabeled data. Target sampling aims to ensure that the generated pseudo labels have a similar distribution with the testing data, which helps the model to learn a more robust and generalizable representation of the data. When the generated pseudo labels have a different distribution than the testing data, the model may be overfitted to the pseudo-labeled data and perform poorly on the test data. As shown in Table 1, using target sampling significantly increased the performance of the model by 11% on ROC-AUC and 115% on PR-AUC, showing the effectiveness of this strategy in improving the performance of MMAPLE.

## 7 Conclusion

In this study, we present MMAPLE, a highly effective deep learning framework, designed to address the challenges of data scarcity and OOD problems encountered when applying machine learning in understudied biological domains when transfer learning is less effective. Through extensive evaluations, we have demonstrated the exceptional capabilities of MMAPLE in exploring the unlabeled data space and facilitating knowledge transfer from one chemical space to another. Using MMAPLE, we successfully predicted and experimentally validated novel interactions between microbiome metabolites and human proteins, thereby shedding light on the intricate interplay between these components. Notably, our framework does not rely on a specific model and can accommodate various deep learning architectures tailored to specific biological tasks. Thus, MMAPLE serves as a versatile and robust framework for investigating a wide range of understudied biological problems.

MMAPLE shows potential for improvement in several key areas. Firstly, the current implementation of MMAPLE lacks the ability to estimate the uncertainty associated with pseudo labels. By incorporating an accurate uncertainty quantification mechanism, it becomes possible to select high-confidence pseudo labels during training, therefore reducing the impact of noise. Secondly, the process of sampling pseudo labels in a vast and imbalanced chemical-protein interaction space proves time-consuming, particularly when aiming to achieve the desired positive versus negative ratio. The performance of MMAPLE can be further enhanced by employing an unbiased and efficient sampling strategy. Lastly, while MMAPLE has thus far been applied exclusively to classification problems, it would be interesting to explore its extension to regression problems.

## 8 Method

### 8.1 Data sets

#### 8.1.1 Protein sequence pre-training dataset

The Pfam database[46] is a comprehensive collection of protein families, represented by Multiple Sequence Alignments (MSA) and hidden Markov models (HMMs). These HMMs are used to identify proteins from new sequences that are likely to belong to a particular family, based on the presence of conserved regions. It provides a wealth of functional annotations for each protein family. It has been shown that pre-trained distilled MSA representation achieved state-of-the-art performance[24]. We used the pre-trained model from this work directly to acquire the protein embeddings. In brief, protein sequence data were first collected from the Pfam database [46], then these sequences were split into clusters based on the 90% of sequence identity, and a representative sequence was chosen from each cluster. The 40,282,439 sequences in the final dataset, which was used to pre-train a transformer model, included a sizable number of GPCRs. We used the original alignment as well as the conservation scores from each Pfam-A family to generate a list of amino acid triplets from the distilled sequence alignment.

#### 8.1.2 Experiment 1: DTI prediction

**Training/validation data** We used molecular interactions in ChEMBL[23] and HMDB[25] as training and validation data. It contained 298,736 total pairs with 230,811 unique chemicals and 3,084 unique proteins.

**OOD testing data** The annotated DTIs from DrugBank[26] were used as testing data. To simulate an OOD scenario, we removed all chemicals that are structurally similar to drugs in the testing data (Tanimoto coefficient¿0.5), totaling 21,760 pairs including 8,917 unique chemicals and 3,266 proteins.

**Unlabeled data** Incorporating a target sampling strategy that focused on the unexplored area within the target distribution, we excluded already labeled pairs, selecting from the remaining candidate unlabeled pairs of proteins and chemicals. For each chemical, we sampled six proteins, resulting in 53,502 total unlabeled pairs. The detailed data statistics can be found in Figure 1.

#### 8.1.3 Experiment 2: Human MPI prediction

**Training/validation data** The training data for this experiment was sourced from ChEMBL (version 29)[23]. It consisted of 334,668 pairs with 252,712 unique compounds and 5,204 unique proteins, where each pair represented an activity with a single protein as the target.

**OOD testing data** For the testing, we utilized HMDB[25], which provided interactions between metabolites and human enzymes. We randomly sampled 10,000 pairs as the testing data covering 8,921 unique compounds and 2,611 unique proteins.

**Unlabeled data** To create the unlabeled dataset, we considered all the unlabeled chemical-protein pairwise combinations. From the total pairs, we included all unique chemicals and randomly selected two enzymes to associate with each chemical, this resulted in the creation of a sizeable unlabeled dataset, consisting of 44,644 unlabeled samples. The detailed data statistics can be found in Figure 1.

#### 8.1.4 Experiment 3: Microbiome-human MPI prediction

**Training/validation data** For this experiment, the training data consisted of a combination of ChEMBL, HMDB, and NJS16[28] datasets. After removing duplicates and unusable data, the dataset contained a total of 1,667,708 samples including 357,213 unique compounds and 168,517 unique proteins.

**OOD testing dataset** The testing dataset was manually created based on two published works. The first work[29] provided information on interactions between 241 GPCRs and metabolites from simplified human microbiomes (SIHUMIs) consisting of the seven most general bacteria species. The second work[30] involved the screening of gut microbiota metabolomes to identify ligands for various GPCRs. Since this study focused on small molecule metabolites, lipids were excluded, resulting in a total of 162 MPIs, including 17 positive activities.

**Unlabeled data** For the protein side, we included all GPCRs from UniProt[47]. Besides, an equal number of proteins were randomly selected from the Pfam dataset. Chemical samples were the 240 unique metabolites from the NJS16 dataset. Overall, the unlabeled data consisted of 73,238 pairs. The detailed data statistics can be found in Figure 1.

### 8.2 MMAPLE base model

Four state-of-the-art baseline models were employed to evaluate the performance MMAPLE:

- DISAE[24]. Distilled Sequence Alignment Embedding (DISAE) is a method developed by us that includes three major modules: protein language model, chemical structure modeling, and the combination of the above two modules. The protein sequence module uses distilled sequence alignment embedding, leveraging a transformer-based architecture trained on nearly half a million protein domain sequences for generating meaningful protein embeddings. In this work, the transformer is trained on approximately half a million protein domain sequences (described in the Data section), allowing it to learn meaningful embeddings for proteins. This is crucial for predicting protein-ligand interactions in out-of-distribution (OOD) scenarios. The chemical module is a graph isomorphism network (GIN) to obtain chemical features, which are numerical representations of small molecules and capture their chemical properties. Finally, DISAE includes an attentive pooling module that combines the protein and chemical embeddings obtained from the first two modules to produce the final output for predicting MPIs as a binary classification task (i.e., active or inactive). The attentive pooling module uses a cross-attention mechanism to weigh the importance of each protein and chemical embedding, allowing it to focus on the most relevant information when making the prediction. *L_base_*denotes the loss function of the base model, which is a binary cross-entropy loss in this case.
- TransformerCPI[16] Adapted from the transformer architecture, TransformerCPI takes protein sequence as the input to the encoder, and atom sequence as the input to the decoder, and learns the interaction at the last layers. Specifically, the amino acid sequence is embedded with a Word2vec model pre-trained on all human protein sequences in UniProt, and the self-attention layers in the encoder are replaced with a gated convolutional network and output the final presentation of proteins. The atom features of chemicals are learned through graph convolutional network (GCN) by aggregating their neighbor atom features. The interaction features are further obtained by the decoder of transfer, which consists of self-attention layers and feed-forward layers.
- DeepPurpose[17] DeepPurpurpose provides a library for DIT prediction incorporating seven protein encoders and eight compound encoders to learn the protein and compound representations respectively, and eventually feeds the learned embeddings into an MLP decoder to generate predictions. We implemented the best-reported architecture, convolutional neural network (CNN) for both protein and compound feature representation learning, as another base model of MMAPLE.
- BACPI[18] The last baseline model included in this study is the Bidirectional attention neural network for compound–protein interaction (BACPI). Similarly, it consists of chemical representation learning, protein representation learning, and CPI prediction components to combine them. BACPI employs a graph attention network (GAT) for compounds to learn various information of the molecule graphs. For protein, it introduces a CNN module to take the amino acid sequence as input, to learn the local contextual features of protein by using a content window to split the sequences onto overlapping subsequences of amino acids. Finally, the atom structure graphs and residue sequence features are fed into the bidirectional attention neural network to integrate the representations and capture the important regions of compounds and proteins, and the integrated features are used to predict the CPI.

### 8.3 Semi-supervised meta-learning

We adopted a semi-supervised meta-learning paradigm for our model training. Similar to Pseudo labels, there is a pair of teacher models and student models, the teacher model takes unlabeled data as input, and uses the predicted results as pseudo labels for the student model to learn with the combination of labeled and pseudo-labeled data. However, instead of learning from the fixed teacher model, the student constantly sends feedback to the teacher in the format of performance on labeled data, and the teacher keeps updating the pseudo labels on every mini-batch. This strategy could solve the problem of confirmation bias in pseudo-labeling[48]. The illustration of MMAPLE training is shown in Figure 5. Let *T* and *S* denote the teacher model and the student model, *θ_T_* and *θ_S_* denote the corresponding parameters (*θ^′^* and *θ^′^* denote the updated parameters). We use *L* to represent the loss function, and *T* (*x_u_*; *θ_T_*) to stand for the teacher predictions on unlabeled data *x_u_*, similar notations for *S*(*x_u_*; *θ_S_*) and *S*(*x_l_*; *θ^′^*). *CE* denotes the cross-entropy loss.

**Figure 5:**
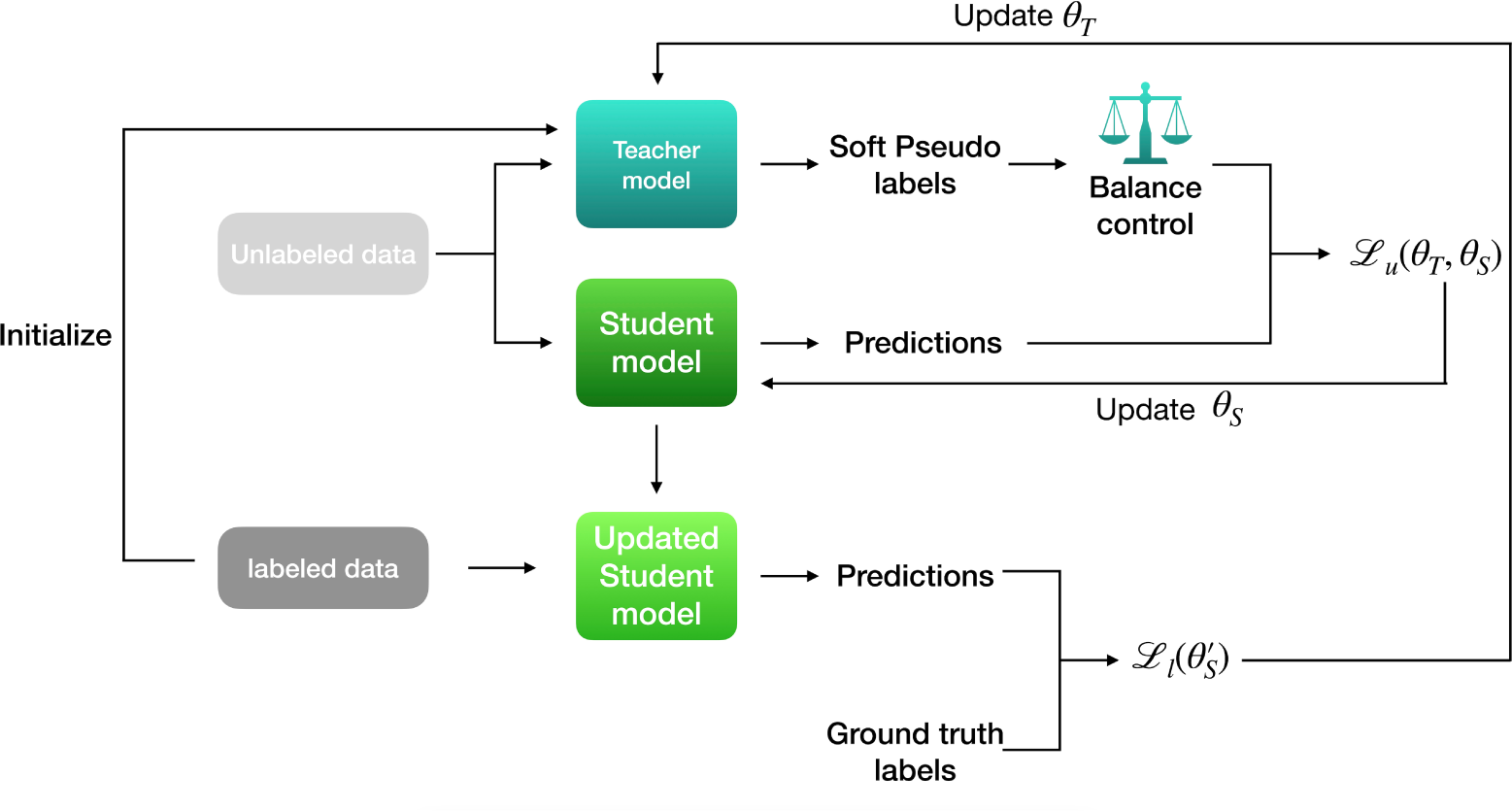
Illustration of MMAPLE training schema. A teacher model generates pseudo labels by predicting a batch of unlabeled data. The pseudo label is further passed to a filter to control the balance ratio of the positive and negative samples (as a hyperparameter). A student model generates the predictions from the same unlabeled data as those used in the teacher model and is updated by minimizing loss function *L_u_*(*θ_T_, θ_S_*) as in equation 3. Then, the updated student model takes a batch of labeled data and generates new predictions that compare with the ground truth labels and minimize loss *L_l_*(*θ^′^*) as equation 6.

### 8.4 Model training

To ensure a fair comparison with the baseline models, both MMAPLE and baseline models were constructed using the same architecture. The detailed training procedure is shown in Algorithm 1.

**The update rule of student** On a batch of unlabeled data *x_u_*, sample *T* (*x_u_*; *θ_T_*) from the teacher’s prediction, and optimize the student model with the objective

where

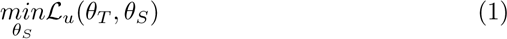

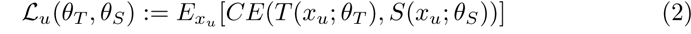

The optimization of each mini-batch is performed as

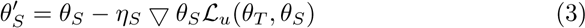

**The update rule of teacher** On a batch of labeled data (*x_l_, y_l_*), and use the students’ update to optimize the teacher model with the objective

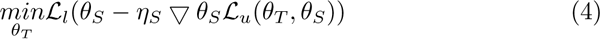

where

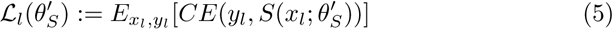

The optimization of each mini-batch is performed as

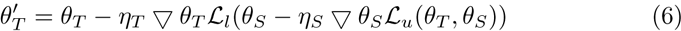

We experimented with both hard labels and soft labels. Due to the superior performance of soft labels to hard labels, the final MMAPLE was trained using the soft label. The methods are described as follows:

**Using soft labels** Because we always treat *θ_S_* as fixed parameters when optimizing Equation 6 and ignore its higher-order dependence on *θ_T_*, the objective is fully differentiable with respect to *θ_T_*when soft pseudo labels are used, i.e., *T* (*x_u_*; *θ_T_*) is the full distribution predicted by the teacher model. This allows us to perform standard back-propagation to obtain the gradient.

Additionally, we incorporated the temperature scaling to soften the teacher model’s predictions [49]. *T* (*x_u_*; *θ_T_*) is the teacher’s output distribution computed by applying softmax over the logits 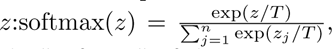 temperature parameter *T* is used to control the ”softness” of the output probabilities. In the implementation, the temperature was tuned by hyperparameter searching.

For the quality control of soft labels, we employed a balance sampler to control the ratio between positive and negative hard labels transferred from soft labels. This will provide a mechanism to dynamically adjust the ratio of positive and negative during training. This ratio served as a crucial parameter to govern the training process, enabling us to strike a balance between the two label categories. Through this approach, we aimed to alleviate bias and imbalance in the dataset.

**Using hard labels** When using hard pseudo labels, we followed the derivative rule proposed in the reference[48], which was a slightly modified version of

REINFORCE applied to obtain the approximated gradient of *L_l_* in Equation6

with respect to *θ_T_*as follows:

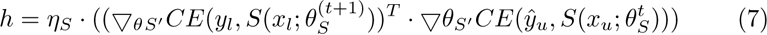

The teacher’s gradient from the student’s feedback:

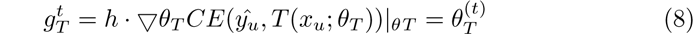

### 8.5 Model evaluation

The model performance was measured using both Receiver Operating Characteristic (ROC) and Precision-Recall (PR) and their corresponding area under the curve (AUC). While ROC is a commonly used metric, it may give an optimistic impression of the model’s performance, particularly when datasets are imbalanced[50]. Therefore, PR is a better metric to evaluate the performance of MMAPLE than ROC. A three-fold cross-validation approach was utilized to ensure the robustness of the model’s performance evaluation. Consistency across evaluations was maintained by using the same folds for all baseline models.

### 8.6 Statistical significance of prediction

In our study, we focused on predicting GPCR genes that interact with TAMO, employing a comprehensive analytical approach to evaluate the statistical significance of each prediction. The prediction scores generated through our model were subjected to Kernel Density Estimation (KDE) from the Python package scipy[51]. KDE is a non-parametric way to estimate the probability density function of our prediction scores. By applying KDE, we were able to calculate the tail probability for each predicted interaction score, which we interpreted as a p-value. This p-value serves as an indicator of the rarity or significance of the predictions within the overall distribution of scores, providing a statistical basis for identifying the most significant GPCR-TAMO interactions. The detailed results of our predictions can be found in Supplemental Table S3.

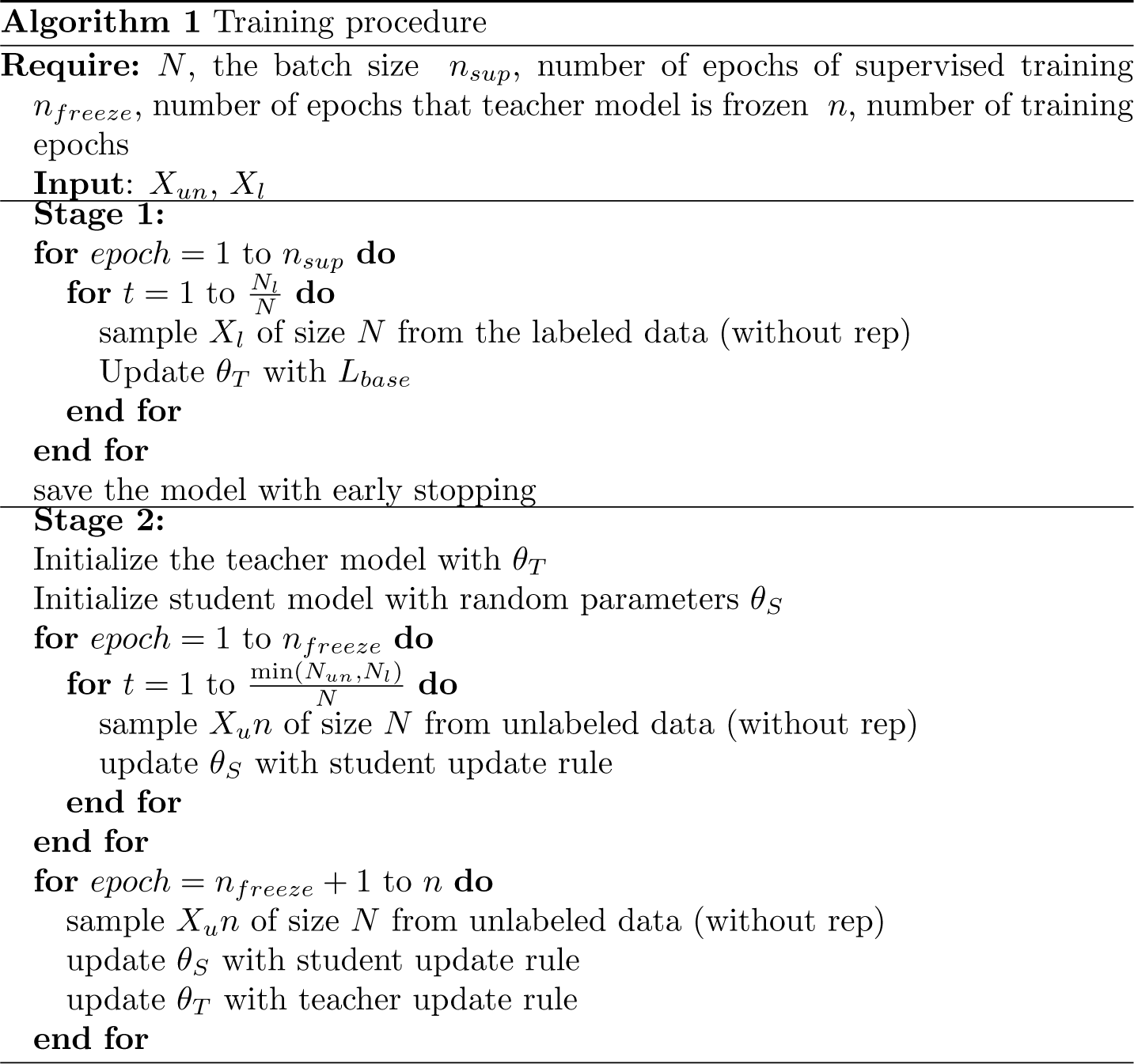

### 8.7 GPCR functional assay

Trimethylamine N-oxide (purity: 95%, molecular weight: 76.12) was purchased from Sigma-Aldrich (MO, USA). GPCR functional assay was performed using the PathHunter *β*-Arrestin assay by Eurofins (CA, USA). The compound activity was analyzed using the CBIS data analysis suite (ChemInnovation, CA).

### 8.8 Protein-ligand docking

AutoDock Vina [36] was applied on TAMO to find the best conformation in the CXCR4 chemokine receptor (PDB ID: 3ODU). The center of the co-crystallized ligand (ligand ID: ITD) in 3ODU was used to define the center of the searching space and 12 Angstrom of extra space was added to the edge of ITD to set up the docking space for TAMO. The binding energies between TAMO and 3ODU were attained in terms of Kcal/mol.

## Data availability

The data used in this study can be accessed at https://doi.org/10.5281/zenodo.10728882 [52].

## Code availability

The source code can be accessed at https://doi.org/10.5281/zenodo.10729017 [53].

## Acknowledgement

This project has been funded with federal funds from the National Institute of General Medical Sciences of the National Institute of Health (R01GM122845), the National Institute on Aging of the National Institute of Health (R01AG057555), and the National Science Foundation (2226183).

## Author Contributions

YW conceived the concept, prepared data, implemented the algorithms, performed the experiments, analyzed data, and wrote the manuscript; Li X performed the experiments, analyzed data, and wrote the manuscript; YL conceived the concept, prepared data, implemented the algorithms, performed the experiments and analyzed data; Lei X conceived and planned the experiments, and wrote the manuscript.

## Competing interests

The authors have declared that no competing interests exist.

## Supplemental materials

**Figure S1:**
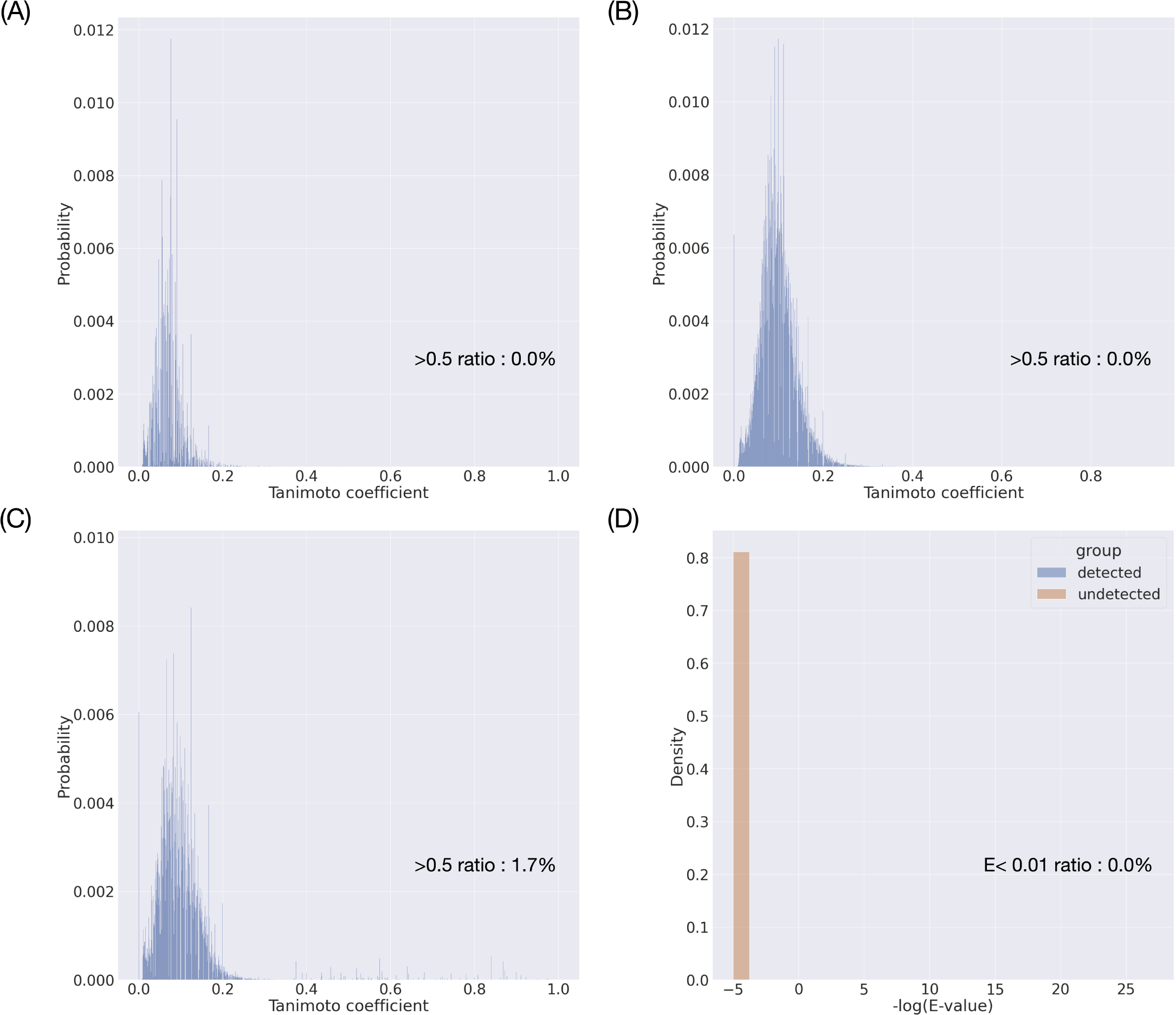
Chemical and protein similarity distribution between training/validation and testing datasets. (A) Chemical similarity distribution of OOD DTI; (B) Chemical similarity distribution of hidden human MPI; and (C, D) Chemical and protein similarity distribution of zero-shot microbiome-human MPI experiment. Chemical similarity is quantified by the Tanimoto coefficient of chemical fingerprints. Protein similarity is measured by the negative logarithm (base 10) of the e-value derived from BLAST[54].

**Figure S2:**
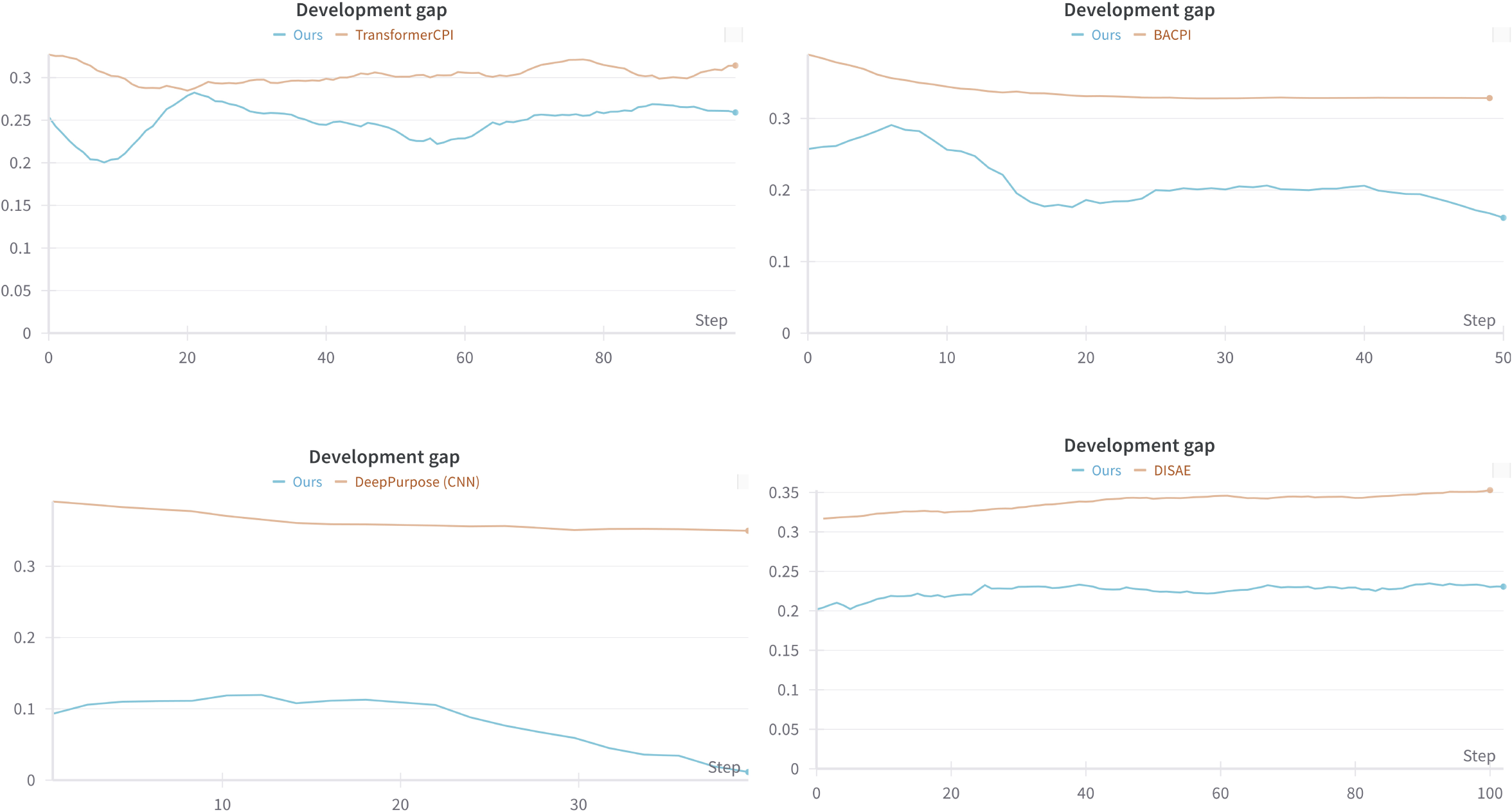
Evaluation of overfitting through the development gap of DTI experiments, calculated as the difference between Validation ROC-AUC and Testing ROC-AUC.

**Figure S3:**
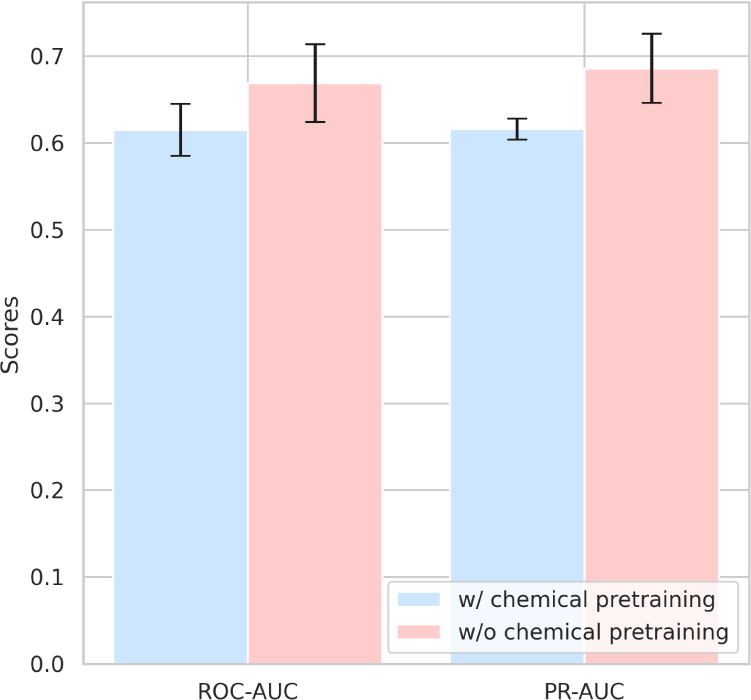
Chemical pre-trained via MolGNN[27] decreases the performance.

**Table S1:** MMAPLE performance over baseline models on ROC-AUC.

|  | Method | DISAE | TransformerCPI | BACPI | DeepPurpose (CNN) |
| --- | --- | --- | --- | --- | --- |
| Human<br>Metabolite-enzyme<br>Interactions | Baseline | 0.669 ± 0.045 | 0.516±0.010 | 0.569±0.044 | 0.622±0.001 |
| | Ours | 0.803 ± 0.046 | 0.606±0.029 | 0.666±0.045 | 0.745±0.054 |
|  | P-val | 0.006 | 0.023 | 0.010 | 0.028 |
|  | Improvement | 20% | 18% | 17% | 20% |
| Drug-target<br>Interaction | Baseline | 0.613±0.007 | 0.604 ± 0.011 | 0.499±0.021 | 0.617 ± 0.011 |
| | Ours | 0.646±0.005 | 0.669 ± 0.031 | 0.640±0.044 | 0.732 ± 0.024 |
|  | P-val | 0.001 | 0.048 | 0.025 | 0.002 |
|  | Improvement | 5% | 11% | 28% | 19% |

**Table S2:**
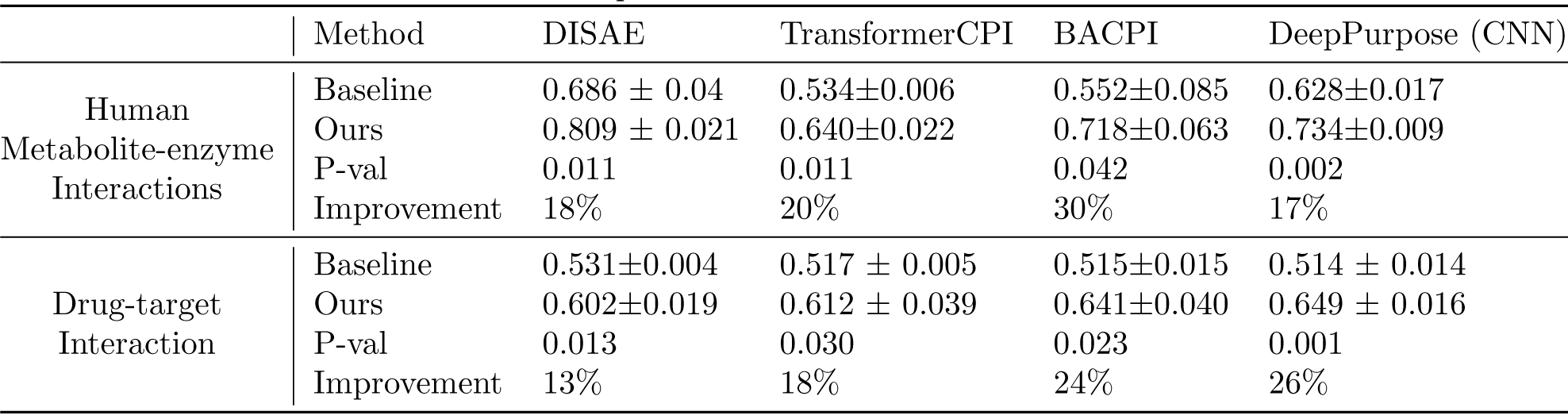
MMAPLE performance over baseline models on PR-AUC.

|  | Method | DISAE | TransformerCPI | BACPI | DeepPurpose (CNN) |
| --- | --- | --- | --- | --- | --- |
| Human<br>Metabolite-enzyme<br>Interactions | Baseline | 0.686 ± 0.04 | 0.534±0.006 | 0.552±0.085 | 0.628±0.017 |
| | Ours | 0.809 ± 0.021 | 0.640±0.022 | 0.718±0.063 | 0.734±0.009 |
|  | P-val | 0.011 | 0.011 | 0.042 | 0.002 |
|  | Improvement | 18% | 20% | 30% | 17% |
| Drug-target<br>Interaction | Baseline | 0.531±0.004 | 0.517 ± 0.005 | 0.515±0.015 | 0.514 ± 0.014 |
| | Ours | 0.602±0.019 | 0.612 ± 0.039 | 0.641±0.040 | 0.649 ± 0.016 |
|  | P-val | 0.013 | 0.030 | 0.023 | 0.001 |
|  | Improvement | 13% | 18% | 24% | 26% |

**Table S3:** Predicted top-ranked G-protein coupled receptor (GPCR) genes that interact with TAMO. p-value stands for the tail probability from Kernel density estimation.

| Rank | Gene | P-val | Rank | Gene | P-val | Rank | Gene | P-val |
| --- | --- | --- | --- | --- | --- | --- | --- | --- |
| 1 | GLP1R | 9.67E-08 | 32 | HRH4 | 8.37E-05 | 63 | TAS2R14 | 1.36E-03 |
| 2 | GIPR | 4.83E-07 | 33 | PTAFR | 8.91E-05 | 64 | GPR151 | 1.39E-03 |
| 3 | CXCR4 | 1.16E-06 | 34 | FFAR4 | 9.86E-05 | 65 | TAS2R39 | 1.77E-03 |
| 4 | CALCRL | 1.61E-06 | 35 | ADORA2B | 1.08E-04 | 66 | GPR37 | 1.78E-03 |
| 5 | GNRHR | 1.93E-06 | 36 | KISS1R | 1.25E-04 | 67 | NPY2R | 1.79E-03 |
| 6 | ADGRA3 | 3.48E-06 | 37 | NPFFR1 | 2.09E-04 | 68 | GRM1 | 1.92E-03 |
| 7 | C3AR1 | 3.80E-06 | 38 | BDKRB1 | 2.58E-04 | 69 | NPY5R | 1.92E-03 |
| 8 | GCGR | 6.67E-06 | 39 | NPSR1 | 2.75E-04 | 70 | CX3CR1 | 2.23E-03 |
| 9 | CCR3 | 1.03E-05 | 40 | TACR2 | 3.05E-04 | 71 | PRLHR | 2.25E-03 |
| 10 | GHRHR | 1.09E-05 | 41 | ADGRE2 | 3.06E-04 | 72 | PTH1R | 2.28E-03 |
| 11 | CCR4 | 1.25E-05 | 42 | HTR4 | 3.71E-04 | 73 | ADGRF1 | 2.61E-03 |
| 12 | CRHR1 | 1.58E-05 | 43 | HTR1B | 3.74E-04 | 74 | ADGRD2 | 2.71E-03 |
| 13 | CALCR | 1.59E-05 | 44 | ADGRG3 | 4.15E-04 | 75 | ADGRG2 | 2.71E-03 |
| 14 | GPR84 | 1.67E-05 | 45 | GRPR | 4.25E-04 | 76 | TACR3 | 2.97E-03 |
| 15 | MLNR | 1.72E-05 | 46 | CCR1 | 4.81E-04 | 77 | ADGRE3 | 3.25E-03 |
| 16 | PTGER4 | 1.98E-05 | 47 | ADGRB3 | 5.05E-04 | 78 | GALR2 | 3.51E-03 |
| 17 | NMUR2 | 2.43E-05 | 48 | OPRL1 | 5.76E-04 | 79 | GHSR | 5.01E-03 |
| 18 | SCTR | 2.90E-05 | 49 | GPR143 | 6.04E-04 | 80 | ADRA2B | 5.45E-03 |
| 19 | VIPR1 | 3.45E-05 | 50 | ADGRG7 | 7.21E-04 | 81 | ADGRA2 | 5.74E-03 |
| 20 | CCKAR | 3.54E-05 | 51 | ACKR2 | 7.23E-04 | 82 | TAS2R8 | 5.87E-03 |
| 21 | MCHR1 | 3.62E-05 | 52 | GPR119 | 7.24E-04 | 83 | HRH1 | 6.88E-03 |
| 22 | CCKBR | 4.09E-05 | 53 | CCR8 | 7.26E-04 | 84 | ADGRF3 | 7.12E-03 |
| 23 | GPR63 | 4.44E-05 | 54 | ADGRF4 | 7.47E-04 | 85 | PTGDR2 | 7.45E-03 |
| 24 | RHO | 4.50E-05 | 55 | OPRM1 | 7.68E-04 | 86 | GLP2R | 9.05E-03 |
| 25 | CCRL2 | 4.73E-05 | 56 | ADGRG4 | 7.87E-04 | 87 | ADRB2 | 1.04E-02 |
| 26 | GPR4 | 6.61E-05 | 57 | NMUR1 | 8.21E-04 | 88 | CYSLTR1 | 1.24E-02 |
| 27 | UTS2R | 6.83E-05 | 58 | S1PR2 | 8.56E-04 | 89 | ADRA1A | 1.26E-02 |
| 28 | HRH3 | 6.90E-05 | 59 | GPR176 | 8.80E-04 | 90 | BDKRB2 | 1.27E-02 |
| 29 | ADORA1 | 7.75E-05 | 60 | ADGRG5 | 8.90E-04 | 91 | GPR183 | 1.27E-02 |
| 30 | HCRTR2 | 7.78E-05 | 61 | CXCR3 | 9.33E-04 | 92 | MC4R | 1.32E-02 |
| 31 | GPBAR1 | 7.92E-05 | 62 | TACR1 | 9.99E-04 | 93 | AVPR1A | 1.47E-02 |

## Notes

### Competing Interest Statement

The authors have declared no competing interest.

### Summary of Updates

Added drug target interaction task; added three baselines including TransformerCPI, BACPI, and DeepPurpose; more post-analysis was also added.

## References

[1] K. G. Hicks, A. A. Cluntun, H. L. Schubert, S. R. Hackett, J. A. Berg, P. G. Leonard, M. A. Ajalla Aleixo, Y. Zhou, A. J. Bott, S. R. Salvatore, et al., “Protein-metabolite interactomics of carbohydrate metabolism reveal regulation of lactate dehydrogenase,” Science, vol. 379, no. 6636, pp. 996– 1003, 2023.

[2] V. Chubukov, L. Gerosa, K. Kochanowski, and U. Sauer, “Coordination of microbial metabolism,” Nature Reviews Microbiology, vol. 12, no. 5, pp. 327–340, 2014.

[3] I. Piazza, K. Kochanowski, V. Cappelletti, T. Fuhrer, E. Noor, U. Sauer, and P. Picotti, “A map of protein-metabolite interactions reveals principles of chemical communication,” Cell, vol. 172, no. 1-2, pp. 358–372, 2018.

[4] M. Groussin, F. Mazel, and E. J. Alm, “Co-evolution and co-speciation of host-gut bacteria systems,” Cell Host & Microbe, vol. 28, no. 1, pp. 12–22, 2020.

[5] M. G. Rooks and W. S. Garrett, “Gut microbiota, metabolites and host immunity,” Nature reviews immunology, vol. 16, no. 6, pp. 341–352, 2016.

[6] G. Bora-Tatar, D. Dayangac-Erden, A. S. Demir, S. Dalkara, K. Yeleķci, and H. Erdem-Yurter, “Molecular modifications on carboxylic acid derivatives as potent histone deacetylase inhibitors: Activity and docking studies,” Bioorganic & medicinal chemistry, vol. 17, no. 14, pp. 5219–5228, 2009.

[7] I. Cho and M. J. Blaser, “The human microbiome: at the interface of health and disease,” Nature Reviews Genetics, vol. 13, no. 4, pp. 260–270, 2012.

[8] J. B. Xavier, V. B. Young, J. Skufca, F. Ginty, T. Testerman, A. T. Pearson, P. Macklin, A. Mitchell, I. Shmulevich, L. Xie, et al., “The cancer microbiome: distinguishing direct and indirect effects requires a systemic view,” Trends in cancer, vol. 6, no. 3, pp. 192–204, 2020.

[9] K. A. Markey, M. R. van den Brink, and J. U. Peled, “Therapeutics targeting the gut microbiome: rigorous pipelines for drug development,” Cell Host & Microbe, vol. 27, no. 2, pp. 169–172, 2020.

[10] T. Cai, L. Xie, S. Zhang, M. Chen, D. He, A. Badkul, Y. Liu, H. K. Namballa, M. Dorogan, W. W. Harding, et al., “End-to-end sequencestructure-function meta-learning predicts genome-wide chemical-protein interactions for dark proteins,” PLOS Computational Biology, vol. 19, no. 1, p. e1010851, 2023.

[11] K. R. Sharma, C. M. Colvis, G. P. Rodgers, and D. M. Sheeley, “Illuminating the druggable genome: Pathways to progress,” Drug Discovery Today, p. 103805, 2023.

[12] G. Kustatscher, T. Collins, A.-C. Gingras, T. Guo, H. Hermjakob, T. Ideker, K. S. Lilley, E. Lundberg, E. M. Marcotte, M. Ralser, et al., “Understudied proteins: opportunities and challenges for functional proteomics,” Nature Methods, vol. 19, no. 7, pp. 774–779, 2022.

[13] T. I. Oprea, C. G. Bologa, S. Brunak, A. Campbell, G. N. Gan, A. Gaulton, S. M. Gomez, R. Guha, A. Hersey, J. Holmes, et al., “Unexplored therapeutic opportunities in the human genome,” Nature reviews Drug discovery, vol. 17, no. 5, pp. 317–332, 2018.

[14] L. Xie, L. Xie, S. L. Kinnings, and P. E. Bourne, “Novel computational approaches to polypharmacology as a means to define responses to individual drugs,” Annual review of pharmacology and toxicology, vol. 52, pp. 361–379, 2012.

[15] A. Sadri, “Is target-based drug discovery efficient? discovery and “offtarget” mechanisms of all drugs,” Journal of Medicinal Chemistry, vol. 66, no. 18, pp. 12651–12677, 2023.

[16] L. Chen, X. Tan, D. Wang, F. Zhong, X. Liu, T. Yang, X. Luo, K. Chen, H. Jiang, and M. Zheng, “Transformercpi: improving compound–protein interaction prediction by sequence-based deep learning with self-attention mechanism and label reversal experiments,” Bioinformatics, vol. 36, no. 16, pp. 4406–4414, 2020.

[17] K. Huang, T. Fu, L. M. Glass, M. Zitnik, C. Xiao, and J. Sun, “Deeppurpose: a deep learning library for drug–target interaction prediction,” Bioinformatics, vol. 36, no. 22-23, pp. 5545–5547, 2020.

[18] M. Li, Z. Lu, Y. Wu, and Y. Li, “Bacpi: a bi-directional attention neural network for compound–protein interaction and binding affinity prediction,” Bioinformatics, vol. 38, no. 7, pp. 1995–2002, 2022.

[19] P. Tossou, C. Wognum, M. Craig, H. Mary, and E. Noutahi, “Real-world molecular out-of-distribution: Specification and investigation,” Journal of Chemical Information and Modeling, vol. 64, no. 3, pp. 697–711, 2024. PMID: 38300258.

[20] Y. Liu, H. Lim, and L. Xie, “Exploration of chemical space with partial labeled noisy student self-training and self-supervised graph embedding,” BMC bioinformatics, vol. 23, no. 3, pp. 1–21, 2022.

[21] B. M. Lake and M. Baroni, “Human-like systematic generalization through a meta-learning neural network,” Nature, vol. 623, no. 7985, pp. 115–121, 2023.

[22] S. Khodadadeh, L. Boloni, and M. Shah, “Unsupervised meta-learning for few-shot image classification,” Advances in neural information processing systems, vol. 32, 2019.

[23] M. Davies, M. Nowotka, G. Papadatos, N. Dedman, A. Gaulton, F. Atkinson, L. Bellis, and J. P. Overington, “Chembl web services: streamlining access to drug discovery data and utilities,” Nucleic acids research, vol. 43, no. W1, pp. W612–W620, 2015.

[24] T. Cai, H. Lim, K. A. Abbu, Y. Qiu, R. Nussinov, and L. Xie, “Msa-regularized protein sequence transformer toward predicting genome-wide chemical-protein interactions: Application to gpcrome deorphanization,” Journal of chemical information and modeling, vol. 61, no. 4, pp. 1570– 1582, 2021.

[25] D. S. Wishart, D. Tzur, C. Knox, R. Eisner, A. C. Guo, N. Young, D. Cheng, K. Jewell, D. Arndt, S. Sawhney, et al., “Hmdb: the human metabolome database,” Nucleic acids research, vol. 35, no. suppl 1, pp. D521–D526, 2007.

[26] D. S. Wishart, Y. D. Feunang, A. C. Guo, E. J. Lo, A. Marcu, J. R. Grant, T. Sajed, D. Johnson, C. Li, Z. Sayeeda, et al., “Drugbank 5.0: a major update to the drugbank database for 2018,” Nucleic acids research, vol. 46, no. D1, pp. D1074–D1082, 2018.

[27] Y. Liu, Y. Wu, X. Shen, and L. Xie, “Covid-19 multi-targeted drug repurposing using few-shot learning,” Frontiers in Bioinformatics, vol. 1, p. 693177, 2021.

[28] J. Sung, S. Kim, J. J. T. Cabatbat, S. Jang, Y.-S. Jin, G. Y. Jung, N. Chia, and P.-J. Kim, “Global metabolic interaction network of the human gut microbiota for context-specific community-scale analysis,” Nature communications, vol. 8, no. 1, pp. 1–12, 2017.

[29] H. Chen, P.-K. Nwe, Y. Yang, C. E. Rosen, A. A. Bielecka, M. Kuchroo, G. W. Cline, A. C. Kruse, A. M. Ring, J. M. Crawford, et al., “A forward chemical genetic screen reveals gut microbiota metabolites that modulate host physiology,” Cell, vol. 177, no. 5, pp. 1217–1231, 2019.

[30] D. A. Colosimo, J. A. Kohn, P. M. Luo, F. J. Piscotta, S. M. Han, A. J. Pickard, A. Rao, J. R. Cross, L. J. Cohen, and S. F. Brady, “Mapping interactions of microbial metabolites with human g-protein-coupled receptors,” Cell host & microbe, vol. 26, no. 2, pp. 273–282, 2019.

[31] R. A. Koeth, Z. Wang, B. S. Levison, J. A. Buffa, E. Org, B. T. Sheehy, E. B. Britt, X. Fu, Y. Wu, L. Li, et al., “Intestinal microbiota metabolism of l-carnitine, a nutrient in red meat, promotes atherosclerosis,” Nature medicine, vol. 19, no. 5, pp. 576–585, 2013.

[32] M. M. Seldin, Y. Meng, H. Qi, W. Zhu, Z. Wang, S. L. Hazen, A. J. Lusis, and D. M. Shih, “Trimethylamine n-oxide promotes vascular inflammation through signaling of mitogen-activated protein kinase and nuclear factor*κ*b,” Journal of the American Heart Association, vol. 5, no. 2, p. e002767, 2016.

[33] S. Yang, X. Li, F. Yang, R. Zhao, X. Pan, J. Liang, L. Tian, X. Li, L. Liu, Y. Xing, et al., “Gut microbiota-dependent marker tmao in promoting cardiovascular disease: inflammation mechanism, clinical prognostic, and potential as a therapeutic target,” Frontiers in pharmacology, vol. 10, p. 1360, 2019.

[34] K. A. Romano, E. I. Vivas, D. Amador-Noguez, and F. E. Rey, “Intestinal microbiota composition modulates choline bioavailability from diet and accumulation of the proatherogenic metabolite trimethylamine-n-oxide,” MBio, vol. 6, no. 2, pp. e02481–14, 2015.

[35] V. E. Brunt, A. G. Casso, R. A. Gioscia-Ryan, Z. J. Sapinsley, B. P. Ziemba, Z. S. Clayton, A. E. Bazzoni, N. S. VanDongen, J. J. Richey, D. A. Hutton, et al., “Gut microbiome-derived metabolite trimethylamine n-oxide induces aortic stiffening and increases systolic blood pressure with aging in mice and humans,” Hypertension, vol. 78, no. 2, pp. 499–511, 2021.

[36] O. Trott and A. J. Olson, “Autodock vina: improving the speed and accuracy of docking with a new scoring function, efficient optimization and multithreading,” Journal of Computational Chemistry, vol. 31, pp. 455– 461, 2010.

[37] M. Kanehisa and S. Goto, “Kegg: kyoto encyclopedia of genes and genomes,” Nucleic acids research, vol. 28, no. 1, pp. 27–30, 2000.

[38] L. Yao, J. Heuser-Baker, O. Herlea-Pana, N. Zhang, L. I. Szweda, T. M. Griffin, and J. Barlic-Dicen, “Deficiency in adipocyte chemokine receptor cxcr4 exacerbates obesity and compromises thermoregulatory responses of brown adipose tissue in a mouse model of diet-induced obesity,” The FASEB Journal, vol. 28, no. 10, p. 4534, 2014.

[39] L. Xu, H. Kitade, Y. Ni, and T. Ota, “Roles of chemokines and chemokine receptors in obesity-associated insulin resistance and nonalcoholic fatty liver disease,” Biomolecules, vol. 5, no. 3, pp. 1563–1579, 2015.

[40] D. L. Costanzo-Garvey, P. T. Pfluger, M. K. Dougherty, J. L. Stock, M. Boehm, O. Chaika, M. R. Fernandez, K. Fisher, R. L. Kortum, E.-G. Hong, et al., “Ksr2 is an essential regulator of amp kinase, energy expenditure, and insulin sensitivity,” Cell metabolism, vol. 10, no. 5, pp. 366–378, 2009.

[41] C.-P. Liang, S. Han, G. Li, I. Tabas, and A. R. Tall, “Impaired mek signaling and serca expression promote er stress and apoptosis in insulin-resistant macrophages and are reversed by exenatide treatment,” Diabetes, vol. 61, no. 10, pp. 2609–2620, 2012.

[42] M. E. Bianchi and R. Mezzapelle, “The chemokine receptor cxcr4 in cell proliferation and tissue regeneration,” Frontiers in Immunology, vol. 11, p. 2109, 2020.

[43] C. Finn, P. Abbeel, and S. Levine, “Model-agnostic meta-learning for fast adaptation of deep networks,” in International conference on machine learning, pp. 1126–1135, PMLR, 2017.

[44] B. Han, Q. Yao, X. Yu, G. Niu, M. Xu, W. Hu, I. Tsang, and M. Sugiyama, “Co-teaching: Robust training of deep neural networks with extremely noisy labels,” in *A*dvances in Neural Information Processing Systems (S. Bengio, H. Wallach, H. Larochelle, K. Grauman, N. Cesa-Bianchi, and R. Garnett, eds.), vol. 31, Curran Associates, Inc., 2018.

[45] C. Thiel, “Classification on soft labels is robust against label noise,” in International Conference on Knowledge-Based and Intelligent Information and Engineering Systems, pp. 65–73, Springer, 2008.

[46] S. El-Gebali, J. Mistry, A. Bateman, S. R. Eddy, A. Luciani, S. C. Potter, M. Qureshi, L. J. Richardson, G. A. Salazar, A. Smart, et al., “The pfam protein families database in 2019,” Nucleic acids research, vol. 47, no. D1, pp. D427–D432, 2019.

[47] “Uniprot: the universal protein knowledgebase in 2023,” Nucleic Acids Research, vol. 51, no. D1, pp. D523–D531, 2023.

[48] H. Pham, Z. Dai, Q. Xie, and Q. V. Le, “Meta pseudo labels,” in Proceedings of the IEEE/CVF Conference on Computer Vision and Pattern Recognition, pp. 11557–11568, 2021.

[49] G. Hinton, O. Vinyals, and J. Dean, “Distilling the knowledge in a neural network,” arXiv preprint arXiv:1503.02531, 2015.

[50] T. Saito and M. Rehmsmeier, “The precision-recall plot is more informative than the roc plot when evaluating binary classifiers on imbalanced datasets,” PloS one, vol. 10, no. 3, p. e0118432, 2015.

[51] P. Virtanen, R. Gommers, T. E. Oliphant, M. Haberland, T. Reddy, D. Cournapeau, E. Burovski, P. Peterson, W. Weckesser, J. Bright, et al., “Scipy 1.0: fundamental algorithms for scientific computing in python,” Nature methods, vol. 17, no. 3, pp. 261–272, 2020.

[52] Y. Wu, “Mmaple dataset v1.0,” Mar. 2024.

[53] Y. Wu, “Xieresearchgroup/mmaple: V1.0,” Feb. 2024.

[54] S. F. Altschul, W. Gish, W. Miller, E. W. Myers, and D. J. Lipman, “Basic local alignment search tool,” Journal of molecular biology, vol. 215, no. 3, pp. 403–410, 1990.

